# Abdominal single-step quantitative susceptibility mapping with spherical mean value filter and structure prior-based regularization

**DOI:** 10.1101/2020.07.15.201327

**Authors:** Anton Abyzov, Bernard E. Van Beers, Philippe Garteiser

**Author notes:** This work was financed in part by France Life Imaging (grant ANR-11-INBS-0006) and by Bpifrance (grant PSPC HECAM). Asterisk indicates corresponding author.

## Abstract

Abdominal quantitative susceptibility mapping (QSM), especially in small animals, is challenging because of respiratory motion and blood flow that, in addition to noise, deteriorate the quality of the input data. Efficient artefact suppression in QSM reconstruction is crucial in these conditions. Single-step QSM algorithms combine background field removal and magnetic field-to-susceptibility inverse problem regularization in a single optimization equation. Here, we propose a single-step QSM algorithm that uses spherical mean value kernels of different radii for background field removal and structure prior (consistency with magnitude image) with *L*_1_ norm for regularization. The optimization problem is solved using the split-Bregman method on the graphic processor unit. The method was compared with previously reported single-step methods: a method using discrete Laplacian instead of spherical mean value kernels, a method using total variational penalty instead of structure prior, and a method using *L*_2_ norm for structure prior. With the proposed method relative to the previous ones, a numerical susceptibility phantom was reconstructed more precisely. In living mice, susceptibility maps with more homogeneous liver, higher contrast between liver and blood vessels, and well-preserved structural details were obtained. In patients, susceptibility maps with more homogeneous subcutaneous fat and higher contrast between subcutaneous fat and liver were obtained. These results show the potential of the proposed single-step method for abdominal QSM in small animals and humans.

## I. INTRODUCTION

Quantitative susceptibility mapping (QSM) is a MRI method that reconstructs the magnetic susceptibility distribution in a tissue [1]. It is currently mostly used in brain imaging, but applications in the body, especially in the liver, start to appear [2]. For instance, QSM has been used as a biomarker for assessing hepatic iron overload [3] and has been proposed to differentiate between patients with simple steatosis and non-alcoholic steatohepatitis (NASH) [4].

Magnetic susceptibility measurements are also a promising way of evaluating the concentration of iron oxide nanoparticles, when used as contrast agents. These nanoparticles have multiple applications in molecular and cellular imaging, especially in studies of macrophage activity [5], [6], and have been proposed to differentiate between NASH and simple steatosis [7]. In contrast to *T*_1_, *T*_2_ and *T*_2_* relaxometry, QSM is not affected by nanoparticle clustering [8], [9], and is less dependent on imaging parameters [10]. The use of QSM for the quantification of iron oxide nanoparticles in small animals has been recently proposed because of the large dynamic range of the method in terms of contrast agent concentration and because of its immunity to susceptibility gradients near organ borders [11].

QSM in living small animals remains challenging, and is more often used in the brain than in the body [12], [13]. QSM of abdominal organs is prone to problems arising from strong spatial variations in tissue susceptibility, chemical shift artifacts related to presence of fat [11], the generally low signal to noise ratio available with current instruments and the short *T*_2_* of liver. Abdominal QSM is further compounded with motion artifacts from respiration, heart contractions and blood flow [14]. Therefore, it is important to minimize errors that result from imperfections of QSM reconstruction algorithms.

The classical QSM reconstruction workflow includes magnetic field assessment from phase data at multiple echo times, background field removal and dipole inversion [1], [15]. The magnetic field can be assessed by analyzing the rate of accumulation of phase with respect to echo time. This process is linear and is made more complex because phase wraps are observed with typical acquisition conditions, and because several proton species (water and fat in the liver) are present in some tissues. Methods such as simultaneous phase unwrapping and removal of chemical shifts (SPURS) [16] or iterative decomposition of water and fat with echo asymmetry (IDEAL) [17], [18] can be proposed.

The magnetic field may be decomposed into an internal field and a background field, both being defined over the complete spatial domain. The background field is the component of the total magnetic field that arises because of susceptibility sources located outside the arbitrary selected reconstruction region, while the internal field results from the susceptibility distribution within this region. Practically, air-tissue interfaces, heterogeneity of the static magnetic field in the scanner and macroscopic currents in the MRI shim coils [19], all occurring outside the reconstructed region, distort the field inside the reconstructed region because of the nonlocal convolution operator and the infinite spatial extent of the dipole kernel.

The final objective of QSM is the determination of the susceptibility distribution within the reconstructed region, hence contributions from the background field need to be carefully handled in the reconstruction. Some susceptibility reconstruction methods include a step of background field removal before dipole inversion [19]. Leporq et al implemented liver susceptibility reconstruction [4] that performed the projection onto dipole field background field removal [20]. More recently, the so-called single step approach has been proposed, wherein the background field removal and dipole inversion steps (see below) are combined together [1], reducing artefacts by avoiding error propagation from one step to another [15]. In a recently published single-step QSM algorithm [15], the background field removal is performed using spherical mean value (SMV) kernels of multiple radii: large radius SMV kernels have better averaging properties, whereas small radius SMV kernels can operate closer to the reconstructed region border, reducing the reconstructed region erosion. Another option is to handle the presence of background field by using the preconditioner in the QSM reconstruction, as in the recently proposed total field inversion method by Liu et al [21].

The dipole inversion problem is an ill-posed one, because the measured field inhomogeneity contains incomplete and corrupted information about underlying susceptibility reconstruction [1], which results in artefacts in reconstructed susceptibility maps. An a priori information (prior) about the susceptibility map may minimize these artefacts by imposing the regularization constraint. The structure prior favors susceptibility maps which structure matches the already known structure (morphology) of the tissue. In the so-called morphology-enabled dipole inversion (MEDI) approach, the regularization is performed by penalizing susceptibility distributions where a gradient in the susceptibility is not accompanied by a strong gradient in the magnitude image (that is usually *T*_2_*-weighted). This MEDI approach has been shown to be very efficient for artefact removal [22], [23]. The *L*_1_ norm (sum of absolute values of all elements) outperforms the *L*_2_ norm (squared root of sum of squares of all elements) in terms of susceptibility artefact removal [24].

In this work, we propose a new single-step QSM algorithm that combines the use of SMV kernels of different radii, regularization using *L*_1_ norm, and magnitude-based structure prior. We compare the performance of this QSM algorithm to that of several recently developed single-step QSM algorithms in a numerical phantom, in the liver of living mice injected with ultrasmall superparamagnetic iron oxide nanoparticles (USPIOs) and in patients with non-alcoholic fatty liver disease.

## II. METHODS

We propose using “multiSMV SS-MEDI”, a single-step MEDI reconstruction involving *L*_1_ norm in the regularization term and a combination of SMV kernels of different radii to remove the background field:

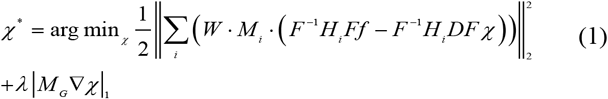

where *H_i_* is the SMV kernel in *k*-space, *D* is the dipole kernel in *k*-space, *W* is the weighting matrix, a diagonal matrix with elements that are equal to the inverse of the standard deviation of the estimated field map [25], *M_G_* is the MEDI structure prior mask derived from the gradient of the magnitude image as proposed by Liu et al [26], *f* is the relative difference field. *M_i_* is the binary trustable region mask for the SMV operator *h_i_. F*^-1^ and *F* are the inverse and direct Fourier transforms, respectively. The extent to which the smoothness in reconstructed susceptibility map is enforced by the regularization term *M_G_∇χ* is controlled by the regularization parameter *λ*. The detailed theoretical background for this equation is provided in the supplementary materials. We compared the proposed method to three single-step reconstruction methods:

1. Discrete Laplacian SS-MEDI, a single-step MEDI reconstruction method that was implemented originally with *L*_2_ norm in the regularization term and a discrete Laplace operator to remove the background field [27]. We use here the *L*_1_ norm instead for the direct comparison with the proposed method, the only difference being the use of discrete Laplacian instead of SMV operators.

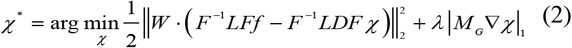
2. multiSMV SS-TV, a single-step method using total variation (TV) regularization with *L*_1_ norm and a combination of SMV kernels of different radii to remove the background field [15]. This method differs from our method by the absence of the structure prior mask *M_G_*. Therefore, all gradients in the reconstructed susceptibility map are minimized (hence the name ‘total variation’).

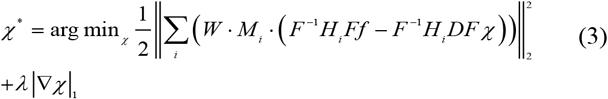
3. MultiSMV SS-MGL2, a single-step structure prior-based reconstruction, involving a combination of SMV kernels of different radii to remove the background field [28]. The difference from the proposed method is the use of *L*_2_ instead of *L*_1_ norm for the regularization term *M_G_*∇*χ*.

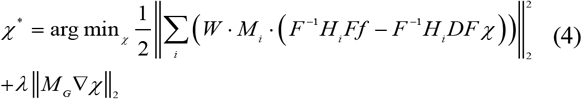

We reconstructed susceptibility maps by finding the minimum of (1) using the ‘split-Bregman’ algorithm [29] proposed and implemented for QSM by Bilgic et al [30]. SMV kernels of radius sizes of 1, 2, 3, 4 and 5 voxels were used. Details of the algorithm implementation with background field removal using SMV kernels can be found in the supplementary materials. Calculations were performed in MATLAB R2017a on a workstation equipped with Intel Core i7 CPU (2.9 GHz, 4 physical cores), and Nvidia Quadro M2200M graphics processing unit (GPU) with 4GB memory on which QSM reconstructions were performed.

### Numerical phantom simulations

The numerical phantom consisted of five cylinders parallel to the magnetic field arranged in a regular pentagon (Fig. 1). The image dimensions were 128 x 128 x 128 voxels. Inside each cylinder there was an ‘insert’ consisting of two superposed smaller cylinders parallel to the outside cylinder. All 5 cylinders were placed in a cylindrical virtual ‘gel’. Uniform susceptibility values of 0, 0.5, 1, 3 and 6 ppm were assigned to the five cylinders, the ‘inserts’ and the ‘gel’ had zero susceptibility, and the susceptibility outside the gel was 9 ppm to simulate the background field created by air around the phantom. Gaussian smoothing using a kernel with diameter of 5 voxels and standard deviation of 0.75 voxel was applied to the susceptibility map to mimic soft borders between cylinders and gel that would appear on real MR images because of partial volume effects. The susceptibilities of the cylinders after Gaussian smoothing were, respectively, 0, 0.444, 0.888, 2.664 and 5.326 ppm.

**Fig. 1.**
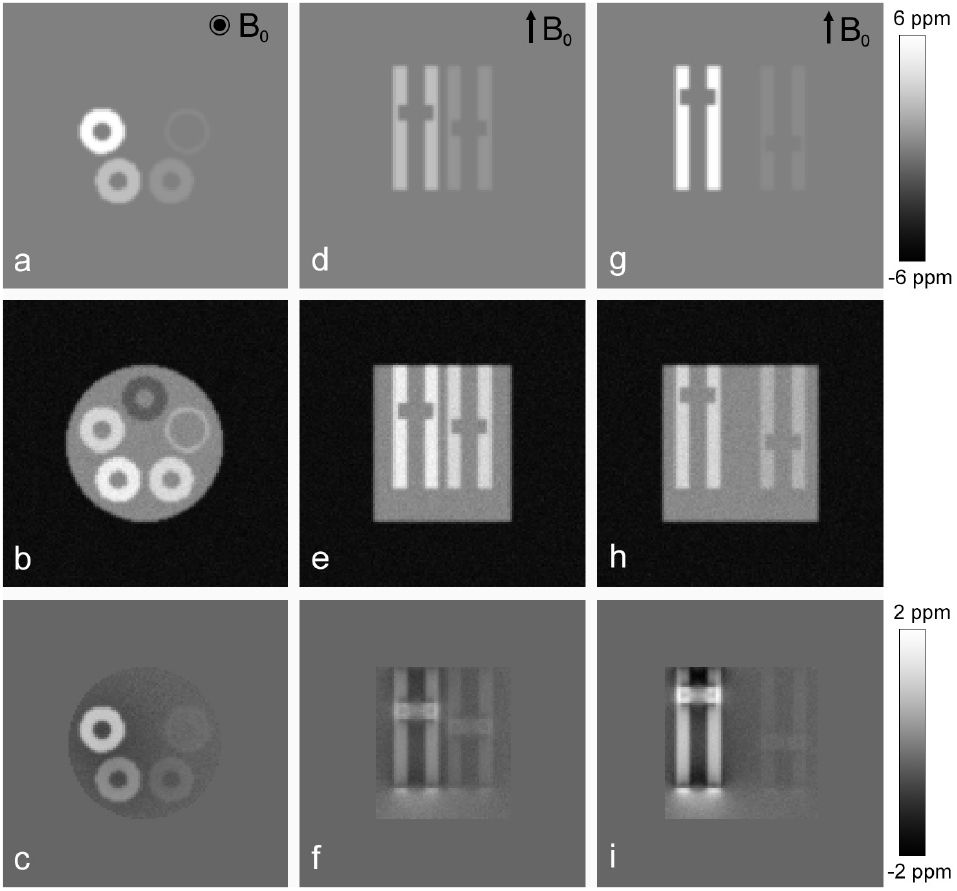
First column: simulated susceptibility map (a), magnitude (b) and relative difference field (c) at a central section in the transverse plane of the numerical phantom. Second and third column: susceptibility map (d,g), magnitude (e,h) and relative difference field (f,i) for two sections in the coronal plane. Two inserts (a thinner and a thicker one) of cylindric shape are seen inside each of the five cylinders.

The relative difference field map was generated from the original susceptibility map by a convolution with a dipole kernel, and was used to obtain phase images for 5 echo times (8.1, 250, 500, 750 and 1000 μs). Corresponding magnitude images were generated using a *T*_2_* exponential decay model, with initial magnitudes of 1·10^4^, 2·10^4^, 2.5·10^4^, 3·10^4^, and 3·10^4^ and *T*_2_* constants of 10, 6.67, 5, 2.5, and 1.43 ms respectively for each of the five cylinders. The ‘gel’ and ‘inserts’ had initial magnitude of 1.5·10^4^ and *T*_2_* constant of 10 ms. The magnitude outside the ‘gel’ was zero. The same Gaussian smoothing as for the susceptibility map was applied to the magnitude images. A complex Gaussian noise with standard deviation of 1000 was added to the complex image, giving a maximal signal-to-noise ratio (SNR) of 30. To estimate the relative difference field, all phase images were unwrapped together using a four-dimensional (4D) version of the phase quality guided path following method [31], [32], the fourth dimension being the echo time. The relative difference field was fitted with a linear least-squares algorithm with echo magnitude weighting [25]. The weighting matrix *W* in the data fidelity term of QSM reconstruction equation was calculated as the inverse of the variance of the fitted relative difference field [25] divided by the mean value of non-zero voxels.

For all QSM methods that used the *L*_1_ norm (discrete Laplacian SS-MEDI, closed-form multiSMV SS-TV, multiSMV SS-MEDI), the stopping criterion for iterations was the relative change in the solution norm becoming smaller than 0.5%. For all methods, except the closed-form multiSMV SS-TV, the MATLAB built-in preconditioned conjugate gradients solver was used with the desired tolerance of 0.005. Regularization parameters (μ and λ) were chosen to minimize the root-mean-square-error (RMSE) of calculated versus original susceptibility. For the closed-form multiSMV SS-TV method, augmented Lagrangian parameters were set to be equal, μ_1_ = μ_2_. For discrete Laplacian and multiSMV SS-MEDI methods that use the ‘split-Bregman’ solver, the μ parameter was chosen to minimize the RMSE in the corresponding *L*_2_ norm problem, as was previously suggested [30]. Optimal parameter values are reported in the supplementary materials.

The magnitude image for MEDI regularization was calculated as square root of the sum of squares of signals at all echo times. The zero referencing was performed by removing the average susceptibility calculated inside the cylinder with initial zero susceptibility from the reconstructed susceptibility map. To evaluate the performance of each method we calculated the RMSE for the whole phantom and individually for the voxels inside each cylinder (excluding the ‘insert’). RMSE values were normalized relative to the original susceptibility, except for the cylinder with zero susceptibility.

### Magnetic resonance imaging in living mice injected with USPIO nanoparticles

Three male C57BL6/J mice were purchased from Envigo (Gannat, France) and housed in the animal facility of the University of Paris (agreement number: C75-18-01) in a temperature-controlled room, with a 12-h light-dark cycle. The mice were 12 week old when the experiments started and stayed in the facility for at least 1 week before MRI. Diet and water were available ad libitum. The experiments were performed according to the procedures approved by our institutional animal care and ethical committee (Apafis #14088). The mice were injected with USPIOs (P01240, CheMatech, Dijon, France) into the tail vein. The injected dose was 100 μmol of Fe (5.6 mg Fe) per kg of mouse mass. Twenty-four hours after USPIO injection, the mice were anesthetized by inhalation of 3% isoflurane before MRI. The body temperature inside the MRI magnet was maintained at 37°C using warm water circulation, and a pressure sensor was used to monitor the respiratory cycle. The MRI data were collected on a Bruker Biospec 7.0T/30 cm USR horizontal magnet (Bruker, Ettlingen, Germany) equipped with the B-GA20S HP shielded gradient set (300 mT/m, 1361 T/m/s) and a ^1^H transmit-receive quadrature coil with 40 mm inner diameter. Nine ultrashort TE 3D sequences were performed with echo times of 0.008, 0.523, 1.045, 1.503, 1.991, 2.468, 2.948, 3.458 and 3.926 ms without respiratory triggering. The other sequence parameters were: TR = 7 ms, non-selective block pulse, flip angle = 5°, field of view = 40 x 40 x 40 mm, matrix size 128 x 128 x 128 (spatial resolution = 0.31 x 0.31 x 0.31 mm), no averaging, bandwidth = 100 kHz, and total acquisition time = 6 min x 9 echoes = 54 min. Magnitude and phase images acquired at different echo times were co-registered: first, the transformation that mutually aligns magnitude images was calculated, and then this transformation was applied to magnitude and phase images.

The trustable region for QSM reconstruction was defined inside the mouse body between two transverse slices, one of which located between the heart and the liver, and the other splitting the right kidney horizontally in two equal parts. Voxels corresponding to the lungs were manually removed to improve reconstruction quality. Phase unwrapping was performed using the 4D algorithm in the same way as for the numerical phantom. The unwrapped images for each echo were further unwrapped using the 3D version of the unwrapping algorithm to better take into account fat signal behavior. Phase unwrapping was unreliable with late echo times, and we used only early echo time images where the average value of the 3D phase derivative variance, used in the unwrapping algorithm, was ≤ 1.5 in the liver ROI (this threshold was chosen empirically based on the appearance of unwrapping artefacts). The relative difference field *f* was calculated taking into account the presence of fat using the T2*-IDEAL approach [33], in which the previously published 9-peak mouse fat spectral model was used [14]. Moreover, the quasi-Newton algorithm in the last step was replaced by the Levenberg-Marquardt algorithm. The weight matrix *W* in the data fidelity term and the magnitude image for MEDI regularization were calculated in the same way as for the numerical phantom.

Stopping criterion for the QSM reconstruction iterations and the conjugate gradients algorithm implementation were the same as for the numerical phantom. The regularization parameter λ was optimized using the L-curve analysis as 4 described by Bilgic et al [30] for all methods. For methods that used ‘split-Bregman’ solver (discrete Laplacian and multiSMV SS-MEDI method), the μ parameter was optimized using the L-curve analysis as described in [30]. For the closed-form multiSMV SS-TV method, the augmented Lagrangian parameters μ1 and μ2 were set to 0.1 as in the original publication [15]. Reconstruction parameters optimized for each mouse are reported in the supplementary materials.

Zero reference mask contained two circular regions with diameter of about 4 voxels placed in spinal muscles on each side of the vertebral column in 30 abdominal transverse slices (9.375 mm) starting from the lung basis. The average susceptibility calculated inside that mask was subtracted from all voxels in the reconstructed susceptibility map. To evaluate the performance of each method we placed on a transverse slice a ROI surrounding four hepatic vessels and a ROI in the liver parenchyma, avoiding vessels and stomach. We compared the mean susceptibility in these ROIs to assess image contrast. To evaluate reconstruction artefacts, we measured the susceptibility homogeneity in the liver by calculating the standard deviation inside the liver ROI, and we compared visually the contours of the hepatic vessels on the magnitude image and the susceptibility map.

### Magnetic resonance imaging in patients

Magnetic resonance imaging data of three patients (62-year old woman, 56-year old man, and 38-year old woman) were taken from a retrospective study of patients with non-alcoholic fatty liver disease (project N°2017-030) for which the local IRB approval was obtained and informed consent was waived because of its retrospective nature. The patients had a liver examination on a Philips Ingenia 3T system (Philips, Best, the Netherlands) with a gradient amplitude of 40 mT/m and a 32-channel phase array body coil and multi-transmit parallel radiofrequency (RF) transmission technology. A 3D mDIXON sequence with parallel imaging and sensitivity encoding (SENSE) was used for imaging with the following parameters: repetition time (TR) 10.3187 ms, flip angle (FA) 5°, bandwidth 2653 Hz/pixel, echo times: 1.1520, 2.3020, 3.4520, 4.6020, 5.7520, 6.9020, 8.0520 and 9.2020 ms. Transverse field of view was 400 x 400 x 220 mm^3^, 400 x 400 x 240 mm^3^, 388 x 388 x 240 mm^3^, and acquisition matrix size was 224 x 224 x 55, 224 x 224 x 60, 224 x 224 x 60 voxels for the three subjects respectively, amounting to a transverse in plane spatial resolution of circa 1.75 mm and equivalent slice thickness of 4 mm, with an acquisition duration of 20 s.

The trustable region for phase unwrapping, relative difference field estimation and QSM reconstruction was segmented automatically with an active contour algorithm and magnitude thresholding. Phase unwrapping was performed with the 3D-SRNCP phase unwrapping algorithm [28]. The water-fat separation with subsequent relative difference field estimation, the weight matrix *W* calculation in the data fidelity term, and the optimization of regularization parameters were performed as described in the previous section. The magnitude image for MEDI regularization was calculated as square root of the sum of squares of signals at even echo times. The optimized reconstruction parameters for each subject are reported. Zero reference mask contained two circular regions in one transverse slice in the spinal muscle. To evaluate the performance of the reconstruction methods, we drew a ROI encompassing the subcutaneous fat and a second ROI in the right liver lobe, avoiding the large vessels. We compared the mean susceptibility of these ROIs to assess image contrast. To assess reconstruction artefacts, we measured the homogeneity of the subcutaneous fat by calculating the susceptibility standard deviation inside the large subcutaneous fat ROI and by comparing the mean susceptibility of two additional smaller ROIs placed in the lateral and the anteroposterior fat regions.

## III. RESULTS

### Numerical phantom

The reconstruction results are reported in Table I and shown in Fig. 2. The reconstruction times were less than 20 seconds with all methods and are reported in the supplementary materials.

**Fig. 2.**
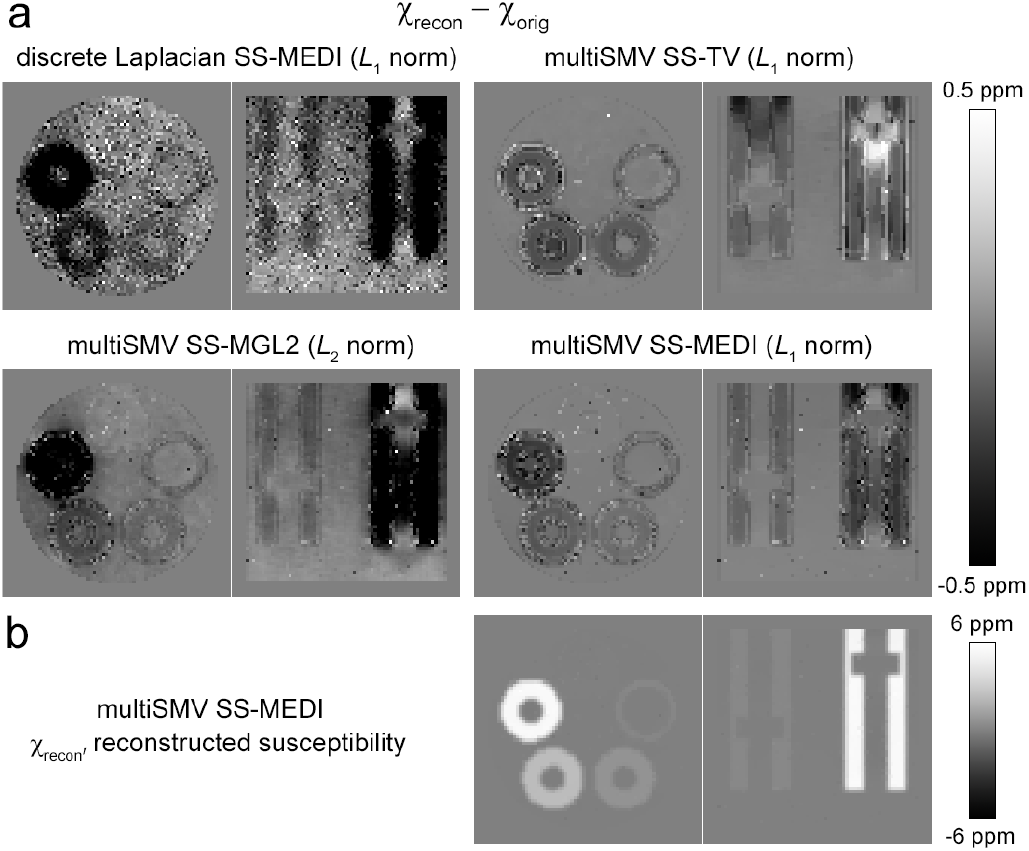
The difference between the reconstructed and the original susceptibility in the numerical phantom using different reconstruction methods (a). A section in the transverse plane and in the coronal plane are shown. The susceptibility map reconstructed with the proposed multiSMV SS-MEDI method (b). The proposed method results in smooth maps with the lowest noise level and minimal differences between reconstructed and original susceptibility maps.

**TABLE I.**
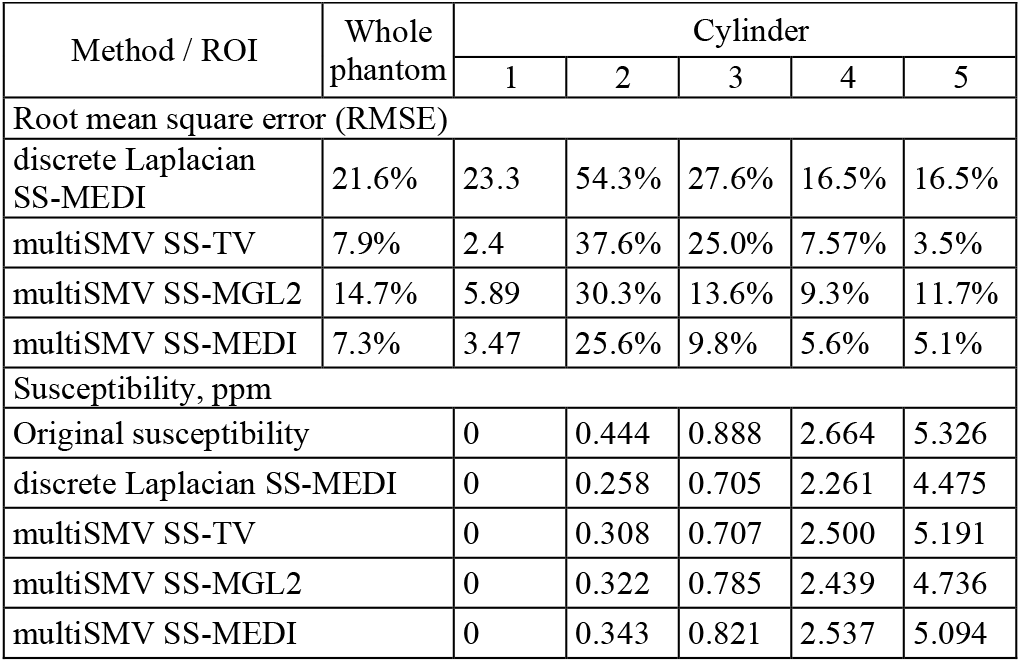
Reconstructed susceptibility in numerical phantom.

The normalized RMSE of calculated relative to original susceptibility was 21.6% with the discrete Laplacian SS-MEDI (highest value), 7.9% with the multiSMV SS-TV, 14.7% with the multiSMV SS-MGL2, and 7.3% with the proposed multiSMV SS-MEDI method (best value). The proposed method was therefore more precise than the other methods.

The susceptibility map reconstructed with the proposed multiSMV SS-MEDI method and difference maps between reconstructed and original susceptibility obtained with all methods are shown in Fig. 2. Despite noisy input data (magnitude and relative difference field map, Fig. 1) the reconstructed susceptibility map obtained with the proposed method was quite smooth. If we take a closer look at the difference maps (Fig. 2), we notice that there was more noise and the susceptibility in all cylinders was significantly underestimated with the discrete Laplacian SS-MEDI method (Table I). In the reconstruction performed with the closed-form SS-TV method, most cylinders had RMSE comparable to that obtained with the proposed method, except for cylinder 3, where the RMSE improvement with the proposed method was substantial (9.8% vs 25%).

In the reconstruction performed with the multiSMV SS-MGL2 method, cylinders 4 and 5 with highest original susceptibility had largely underestimated susceptibility values. In conclusion, the original susceptibility was underestimated with all methods, but to a lesser extent with the proposed method.

### Mice injected with USPIO nanoparticles

Reconstruction results are reported in Table II for the three mice and shown in Fig. 3 for the first mouse. The reconstruction times were less than one minute for all mice and are reported in the supplementary materials.

**TABLE II.** Liver and vessel Susceptibility in mice injected with USPIO.

| # | Method | Susceptibility, ppm |  |  |
| --- | --- | --- | --- | --- |
|  |  | Liver |  | Vessels |
|  |  | Mean | St. dev. | Mean |
| 1 | discrete Laplacian SS-MEDI | 0.523 | 0.085 | 0.373 |
|  | multiSMV SS-TV | 0.245 | 0.135 | 0.030 |
|  | multiSMV SS-MGL2 | 0.574 | 0.065 | 0.376 |
|  | multiSMV SS-MEDI | 0.876 | 0.049 | 0.554 |
| 2 | discrete Laplacian SS-MEDI | 0.588 | 0.062 | 0.371 |
|  | multiSMV SS-TV | 0.158 | 0.083 | 0.009 |
|  | multiSMV SS-MGL2 | 0.547 | 0.058 | 0.275 |
|  | multiSMV SS-MEDI | 0.718 | 0.038 | 0.446 |
| 3 | discrete Laplacian SS-MEDI | 0.146 | 0.046 | -0.105 |
|  | multiSMV SS-TV | 0.292 | 0.250 | 0.034 |
|  | multiSMV SS-MGL2 | 0.383 | 0.054 | 0.226 |
|  | multiSMV SS-MEDI | 0.637 | 0.038 | 0.385 |

**Fig. 3.**
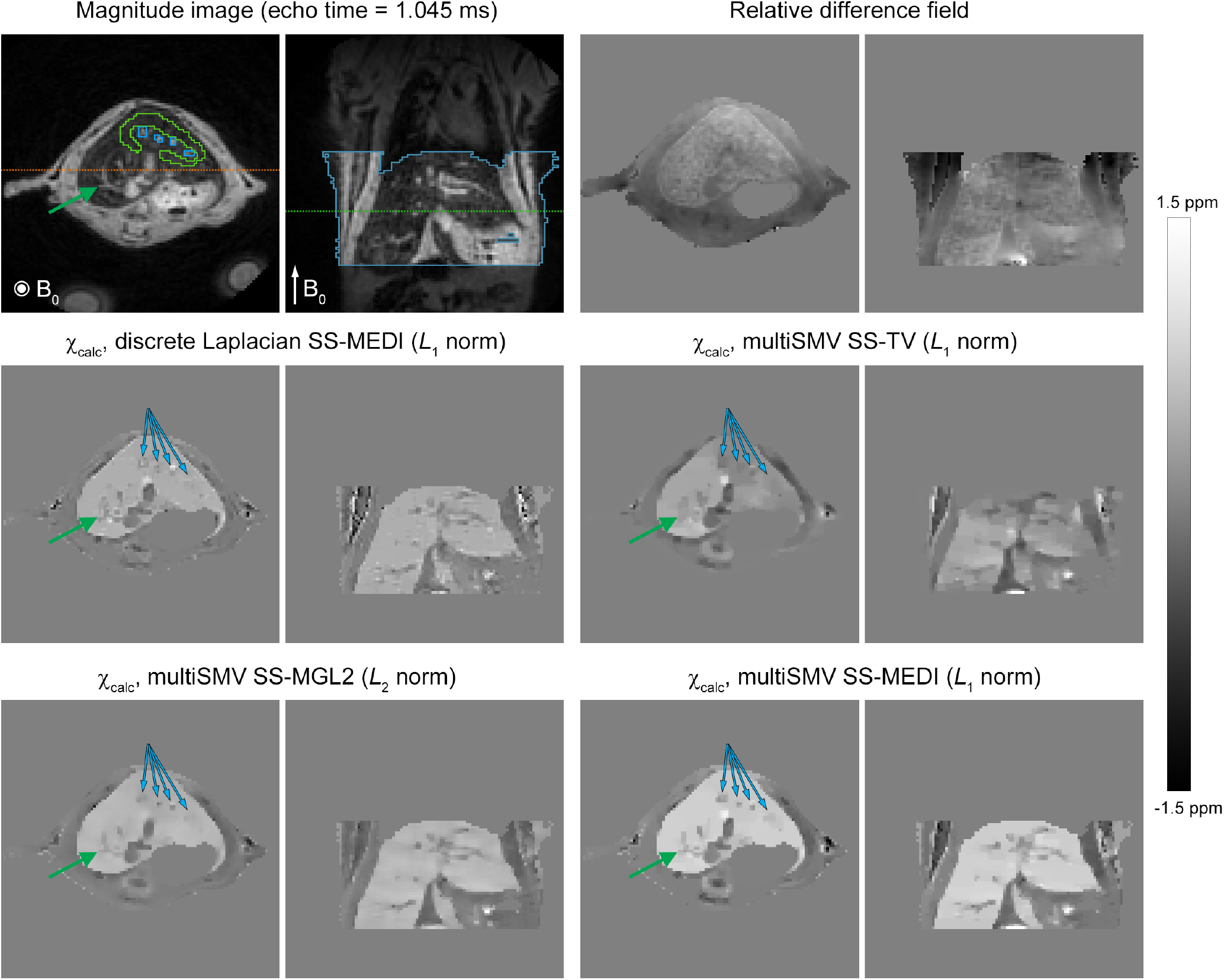
Mouse injected with USPIO nanoparticles. Magnitude, relative difference field and reconstructed susceptibility maps obtained with the different methods, in transverse and coronal planes. The liver appears dark on the magnitude image, and the blood vessels appear bright. Orange dotted line shows the position of the coronal slice in the transverse plane, and green dotted line shows the position of the transverse slice in the coronal plane. Green contour: ROI in the liver in the immediate vicinity of vessels. Blue contours ROI around vessels. On the magnitude image in the coronal plane, blue contour surrounds voxels inside reconstructed region, those outside it are shadowed. Blue and green arrows: positions of small vessels. The scale is shown for the relative difference field and susceptibility images. The proposed multiSMV SS-MEDI method results in smooth maps while still displaying high level of structural details.

With the discrete Laplacian SS-MEDI method, the liver susceptibility was relatively homogeneous (standard deviation 0.046 – 0.085 ppm, Table II). However, the contours of small vessels (blue and green arrows) on the reconstructed susceptibility images showed artefacts. With the multiSMV SS-TV reconstruction, the liver susceptibility was much more heterogeneous (Fig. 3), and its standard deviation was the highest among all reconstructions (0.083 – 0.250 ppm). These heterogeneities, not observed in the magnitude image, resulted from reconstruction artefacts. The vessel contours were lost.

With the multiSMV SS-MGL2 method and with the proposed multiSMV SS-MEDI method, the vessels were quite well delineated. The susceptibility of the liver was more homogeneous with the proposed method than with the multiSMV SS-MGL2 method (standard deviation was 0.038 – 0.049 ppm vs 0.054 – 0.065 ppm). The contrast between liver and vessels was higher with the proposed method (mean susceptibility differences of 0.252-0.322 ppm) than with the other methods (differences of 0.150-0.272 ppm).

We also observed that the mean values of liver susceptibility were higher with the proposed multiSMV SS-MEDI reconstruction (0.637 – 0.876 ppm) than with the other reconstructions (0.146 – 0.588 ppm among all mice and methods).

### Patient imaging

The reconstruction results are reported in Table III for the three patients and shown in Fig. 4 for the first patient. Reconstruction times were less than 250 seconds with the proposed method and less than 130 seconds with other methods (supplementary materials). The liver susceptibility maps were homogeneous with all reconstruction methods, with standard deviations always below 0.02 ppm.

**Figure 4.**
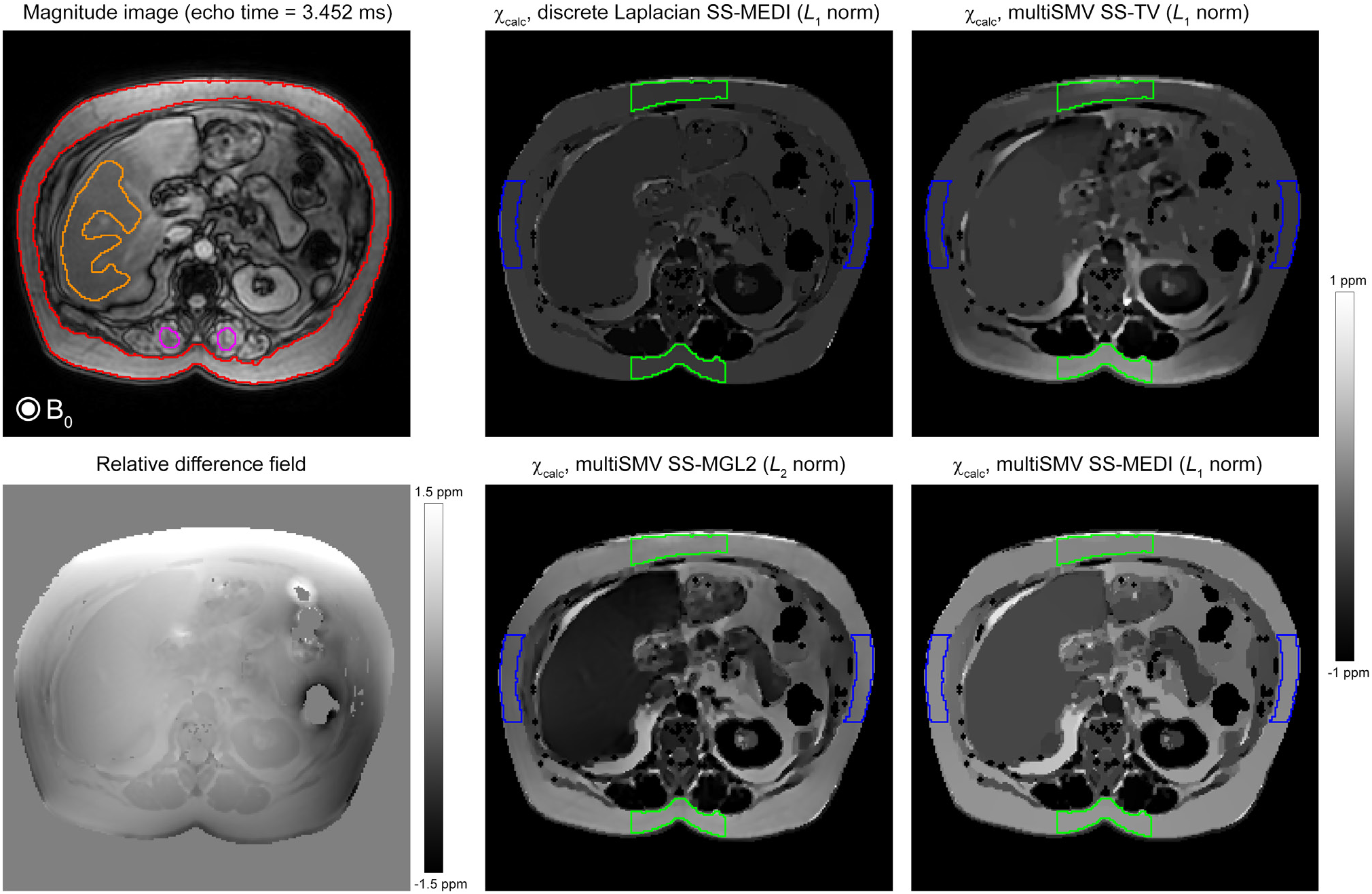
Patient MRI. Magnitude, relative difference field and susceptibility maps for the different reconstruction methods. Red contour: ROI in the subcutaneous fat. Orange contour: ROI in the liver. Magenta contour: ROI in the spinal muscles, the average susceptibility of which was subtracted from the reconstructed susceptibility map for the zero reference. Blue and green contours represent the lateral and the anteroposterior regions respectively inside the subcutaneous fat ROI. The proposed reconstruction proposed multiSMV SS-MEDI method displays high liver to fat contrast and strong homogeneity in subcutaneous fat.

With the discrete Laplacian SS-MEDI reconstruction, the liver and subcutaneous fat had susceptibility values that were very close (mean susceptibility difference of 0.06-0.126 ppm) (Fig. 4 and Table III). With the closed-form multiSMV SS-TV reconstruction, the susceptibility was heterogeneous in subcutaneous fat with the highest standard deviation (0.065 – 0.214 ppm). The liver to fat contrast was also low with the multiSMV SS-TV reconstruction (0.046-0.185 ppm).

**TABLE III.** Susceptibility in patient subcutaneous fat and liver.

| # | Method | Susceptibility, ppm |  |  |  |  |
| --- | --- | --- | --- | --- | --- | --- |
|  |  | Subcutaneous fat ROI |  | Antero-posterior or region | Lateral relative to anteroposterior or region, % | Liver ROI |
|  |  | Mean | St. dev. | Mean | Mean | Mean |
| 1 | discrete Laplacian SS-MEDI | 0.212 | 0.028 | 0.213 | 0.214 (+0.7%) | 0.152 |
|  | multiSMV SS-TV | 0.302 | 0.111 | 0.413 | 0.226 (-45.3%) | 0.224 |
|  | multiSMV SS-MGL2 | 0.491 | 0.069 | 0.547 | 0.407 (-25.7%) | 0.114 |
|  | multiSMV SS-MEDI | 0.545 | 0.040 | 0.551 | 0.537 (-2.4%) | 0.275 |
| 2 | discrete Laplacian SS-MEDI | 0.117 | 0.035 | 0.145 | 0.086 (-40.4%) | 0.028 |
|  | multiSMV SS-TV | 0.297 | 0.214 | 0.458 | 0.130 (-71.7%) | 0.112 |
|  | multiSMV SS-MGL2 | 0.523 | 0.109 | 0.605 | 0.391 (-35.5%) | 0.137 |
|  | multiSMV SS-MEDI | 0.594 | 0.042 | 0.603 | 0.564 (-6.5%) | 0.243 |
| 3 | discrete Laplacian SS-MEDI | 0.16 | 0.017 | 0.164 | 0.153 (-6.3%) | 0.034 |
|  | multiSMV SS-TV | 0.174 | 0.065 | 0.225 | 0.135 (-40.1%) | 0.128 |
|  | multiSMV SS-MGL2 | 0.552 | 0.057 | 0.592 | 0.488 (-17.4%) | 0.096 |
|  | multiSMV SS-MEDI | 0.688 | 0.023 | 0.678 | 0.695 (2.5%) | 0.137 |

Higher contrast was obtained with the multiSMV SS-MGL2 and SS-MEDI reconstructions (mean susceptibility differences of 0.377-0.456 ppm and 0.270-0.551 ppm, respectively). With the proposed multiSMV SS-MEDI method, the susceptibility maps of subcutaneous fat were more homogeneous (standard deviation 0.023 – 0.042 ppm, difference between mean values in anterioposterior and lateral regions 2.5% – 6.5%) than with the multiSMV SS-MGL2 reconstruction (standard deviation 0.057 – 0.109 ppm, difference 17.4% – 35.5%).

## IV. DISCUSSION

The results that we obtained in numerical phantom simulations, in mouse liver enhanced with USPIO nanoparticles and in patient liver showed that the contrast and image quality of susceptibility reconstructions are improved by combining the single-step QSM approach with SMV kernels of different radii, regularization with *L*_1_ norm, and structure prior.

When the background field is removed using the discrete Laplacian instead of SMV kernels, as in the discrete Laplacian SS-MEDI method, reconstructed susceptibility maps were of a bad quality. In numeric phantom simulations, the highest RMSE was obtained using this method, and high levels of noise were observed in the susceptibility maps. In mice, low contrast between liver and vessels was observed, as well as distorted small vessel contours. In patients, low contrast between liver and subcutaneous fat was seen. The discrete Laplacian had a kernel size of only 3 x 3 x 3 voxels and was more prone to noise amplification than larger SMV kernels with better averaging properties. These features reduced the quality of the reconstructed susceptibility map with the discrete Laplacian method.

The absence of structure prior in multiSMV SS-TV method also resulted in QSM reconstructions of lesser quality because the use of TV-regularization penalized all gradients in the reconstructed susceptibility map. In the mouse liver susceptibility maps obtained with the multiSMV SS-TV method the details (such as small vessel contours) were lost. This was likely caused by the lack of structural information at the interfaces between liver and vessels in the relative difference field map (Fig. 3).

With the reconstructions that include structure priors (multiSMV SS-MGL2 and multiSMV SS-MEDI) the previously mentioned problems were minimized. The liver susceptibility in mice and the subcutaneous fat susceptibility in patients were heterogeneous with the multiSMV SS-TV reconstruction, whereas the inclusion of the magnitude image structure prior rendered the subcutaneous fat more homogeneous.

The regularization with the *L*_2_ norm in multiSMV SS-MGL2 resulted in poorer RMSE in numeric phantom simulations, and more heterogeneous susceptibility maps in mouse and patient MRI. The *L*_2_ norm used in multiSMV SS-MGL2 is less performant in structural matching between magnitude and susceptibility maps than the *L*_1_ norm [23], [35]. It has also been reported that the *L*_2_ norm produces susceptibility maps that are less homogeneous because of artefacts, both in simulations and in brain imaging [23], [24].

The use of the *L*_1_ norm in the proposed multiSMV SS-MEDI method allowed us reconstructing the numerical phantom more precisely, better mitigating artefacts, and increasing the homogeneity of the susceptibility maps in mice and patients while maintaining a high degree of contrast detail. This came, however, at the price of a somewhat longer reconstruction time, which remained shorter than 5 minutes per image in all cases.

As we pointed out above, the mean susceptibility values obtained in numerical phantoms, mouse and patient liver with the proposed multiSMV SS-MEDI method were higher than with the other methods. In the numerical phantoms, these higher values were also closer to the original values. We suggest that the liver susceptibility obtained with our method in mouse and patient MRI might be the most exact one, as this would be in agreement with numerical phantom simulations. However, to confirm this suggestion, we would need a reference examination for susceptibility measurements, such as COSMOS QSM [36] or superconducting quantum interference device-based liver susceptometry [37]. This absence of reference method is a limitation of our study.

We also did not probe different USPIO concentrations in mouse liver MRI to evaluate the response of different QSM reconstruction methods to a range of iron oxide nanoparticle concentrations. From the methodology point of view, we should point out that the discrete Laplacian SS-MEDI method used in our comparison was originally implemented with the *L*_2_ norm [27], but we chose to reimplement it with the *L*_1_ norm so that the only difference evaluated would be the background field removal operator. We expect that the combination of *L*_2_ norm and discrete Laplacian, as in the original implementation, would further reduce the method diagnostic performance.

Moreover, it might be useful to compare the single-step QSM methods with methods that address the background field removal differently, as the preconditioned total field inversion QSM [21] that was recently used in small animals [11]. Finally, the scope of our study was limited to background field removal and dipole inversion in the QSM reconstruction workflow. To further optimize QSM for abdominal imaging, future work should be performed to compare magnetic field calculation methods, such as IDEAL and SPURS.

## V. CONCLUSION

We propose a new QSM reconstruction method that combines the single-step dipole inversion approach with SMV kernels of different radii, regularization using the *L*_1_ norm and structure prior. We validated the performance of the proposed method in numerical phantom simulations, and showed that the method allowed reconstructing susceptibility maps with less artefacts, more contrast, homogeneity and structural details than other previously described single-step methods in the upper abdomen of small animals and patients.

## Acknowledgment

The authors thank Dr. Valerie Vilgrain, the coordinator of the clinical study from which patient MRI data were used in this paper.

## Notes

### Competing Interest Statement

The authors have declared no competing interest.

