## Supplementary Material for "Abdominal single-step quantitative susceptibility mapping with spherical mean value filter and structure prior-based regularization"

**SUPPLEMENTARY INFORMATION**

*QSM reconstruction parameters and times*

**Table S1.** Parameters for QSM reconstruction methods.

| Subject | discrete Laplacian SS-MEDI | | multiSMV SS-TV | | multiSMV SS-MGL2 | multiSMV SS-MEDI | |
| --- | --- | --- | --- | --- | --- | --- | --- |
|  | μ | λ | μ_1_ = μ_2_ | λ | λ | μ | λ |
| Numerical  phantom | 6·10^-2^ | 4·10^-2^ | 5·10^-3^ | 2·10^-3^ | 2·10^-1^ | 2·10^-1^ | 8·10^-3^ |
| Mouse 1 | 3.16 | 5.18·10^-2^ | 0.1, fixed | 5.18·10^-3^ | 1.93·10^-1^ | 1.93·10^-1^ | 7.20·10^-3^ |
| Mouse 2 | 7.19·10^-1^ | 3.16·10^-2^ | 0.1, fixed | 5.18·10^-3^ | 1.18·10^-1^ | 1.18·10^-1^ | 7.20·10^-3^ |
| Mouse 3 | 3.16 | 8.48·10^-2^ | 0.1, fixed | 5.18·10^-3^ | 8.48·10^-1^ | 8.48·10^-1^ | 1.93·10^-2^ |
| Patient 1 | 3.16 | 5.18·10^-2^ | 0.1, fixed | 3.16·10^-3^ | 4.39·10^-1^ | 4.39·10^-1^ | 1.93·10^-2^ |
| Patient 2 | 3.16 | 8.48·10^-2^ | 0.1, fixed | 3.16·10^-3^ | 7.20·10^-1^ | 7.20·10^-1^ | 3.16·10^-2^ |
| Patient 3 | 1.93 | 5.18·10^-2^ | 0.1, fixed | 5.18·10^-3^ | 7.20·10^-1^ | 7.20·10^-1^ | 1.18·10^-2^ |

**Table S2.** Reconstruction times for QSM reconstruction methods.

| Subject | discrete Laplacian SS-MEDI | | multiSMV SS-TV | multiSMV SS-MGL2 | | multiSMV SS-MEDI | |
| --- | --- | --- | --- | --- | --- | --- | --- |
|  | Time [s] | Conjugate gradient  iterations | Time [s] | Time [s] | Conjugate gradient  iterations | Time [s] | Conjugate gradient  iterations |
| Numerical  phantom | 12.0 | 92 | 18.8 | 5.0 | 27 | 12.9 | 59 |
| Mouse 1 | 25.2 | 195 | 10.9 | 10.7 | 61 | 37.2 | 181 |
| Mouse 2 | 28.3 | 222 | 12.0 | 12.4 | 72 | 41.1 | 202 |
| Mouse 3 | 25.7 | 184 | 12.0 | 25.6 | 152 | 59.0 | 318 |
| Patient 1 | 129.0 | 441 | 44.4 | 77.0 | 111 | 246.8 | 322 |
| Patient 2 | 129.8 | 374 | 38.0 | 64.1 | 108 | 223.1 | 346 |
| Patient 3 | 100.9 | 346 | 29.0 | 62.1 | 122 | 174.8 | 322 |

*QSM reconstruction: theoretical considerations*

In the presence of an applied static magnetic field, the magnetic susceptibility distribution in a tissue creates a magnetic field heterogeneity. The susceptibility *χ* can be found as a solution of the following minimization problem (called dipole inversion):

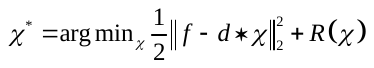
(1)

Here in the first data fidelity term, *f* is the relative difference field (normalized magnetic field inhomogeneity), *d* is the dipole kernel, and *R*(*χ*) is a regularization term (see below). *χ** is the susceptibility map that solves the problem. The convolution operation * in (1) is calculated in the Fourier space as a multiplication:

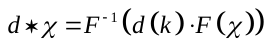
(2)

Here *d*(*k*) is the dipole kernel in the Fourier (**k**) space:

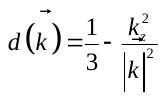
(3)

And *F^-1^* and *F* are the inverse and direct Fourier transforms.

The relative difference field *f* is generated by susceptibility sources inside the reconstructed region, but it also contains the contribution of the background field. Taking into account the susceptibility sources originating outside of the reconstruction volume would be intractable because of problem dimensionality and because of signal lack in large regions of the domain. When decomposing the background field contributions from the total field in a separate, prior step, the reconstructed susceptibility map suffers from error propagation from one step to another [1], [2]. In one-step approaches, the background field is removed directly inside the dipole inversion calculation:

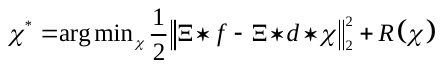
(4)

Here Ξ is a background field removal operator such as:

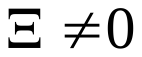
;
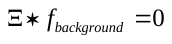
(5)

In this context the main property of the background field is that it is harmonic inside the reconstructed region, hence it is possible to remove it by applying an operator that selectively suppresses harmonic functions [3]. The Laplace operator Δ is one of such operators, but any positive function with radial symmetry and integral equal to one (normalized), also fulfills this condition [4], [5]. In practice, it is implemented as a discrete operator that uses only neighboring voxels to calculate the result and its output suffers from noise. Schweser et al. have proposed to use the spherical mean value (SMV) operator [3]:

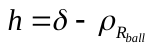
(6)

where *ρ_Rball_* is a ball of a radius *R_ball_* at the center of coordinates, such that:

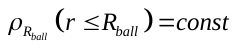
,
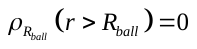
,
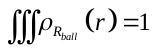
(7)

and *δ* is the delta function at the center of coordinates.

The choice of SMV operator radius is very important, because bigger radius means larger number of voxels averaging and better resilience to error amplification [6], [7] However, the Laplace or SMV operators should be evaluated inside the region in which the relative difference field is calculated. This results in the erosion of the reconstructed region, as the operator cannot be evaluated too close to the region edge. For the discrete Laplace operator, a one voxel-thick layer is eroded. For the SMV operator with *R_ball_* radius, a layer of approximately *R_ball_* voxels near the edge is lost. A combination of SMV operators of different radii has been recently proposed [2], [8] to realize a high-fidelity background field removal while limiting the erosion of the trustable region to one voxel:

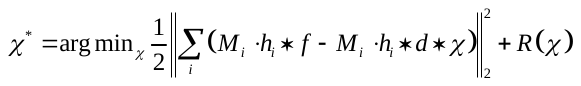
(8)

where *M_i_* is the binary trustable region mask for the SMV operator *h_i_* of radius *i*, that applies to each voxel a SMV kernel of the largest possible radius (small radius for peripheric voxels and larger radius for central voxels).

The equation that we use to calculate the susceptibility in the proposed method is the following:

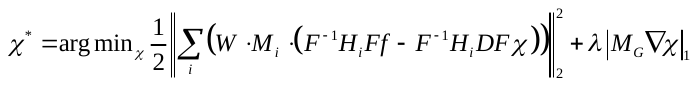
(9)

where *H_i_* is the SMV kernel *h_i_* in *k*-space, *D* is the dipole kernel in *k*-space, *W* is the weighting matrix, a diagonal matrix with elements that are equal to the inverse of the standard deviation of the estimated field map [9], and *F^-1^* and *F* the inverse and direct Fourier transforms, respectively. We replace convolution operators in (8) by the multiplication in the Fourier (*k*) space, according to (2). *F^-1^H_i_DFχ* means the successive application of the direct Fourier transform *F*, multiplication in the *k*-space with the dipole kernel *D*, multiplication in the *k*-space with the SMV kernel *H_i_*, and application of the inverse Fourier transform *F^-1^* to the reconstructed susceptibility map *χ*.

The last term, |*M_G_*∇χ|_1_, is the MEDI regularization term as implemented in [10]. It contains a structure prior mask *M_G_* derived from the gradient of the magnitude image. In this mask, 30% of voxels with highest magnitude image gradient values are arbitrarily set to zero and the others to the value of one, according to the results obtained by Liu et al [11]. By design, this regularization term enforces smoothness in the susceptibility distribution based on image regions where the magnitude is itself smooth. The extent to which the smoothness is enforced is controlled by the regularization parameter *λ*.

*Computational approach*

The minimization of the data fidelity term, as it contains a *L*_2_ norm, is quite straightforward with the conjugate gradient algorithm [12]. However, the minimization of a *L*_1_ norm in MEDI regularization requires a more complex algorithm, such as the ‘split-Bregman’ iterative solver [13] proposed and implemented for the QSM dipole inversion by Bilgic et al [14]. The constrained optimization problem derived from (9):

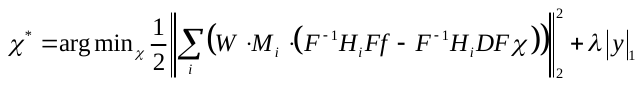
, such that
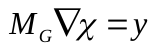
(10)

is relaxed to an unconstrained optimization problem, and solved iteratively with the ‘split-Bregman’ approach:

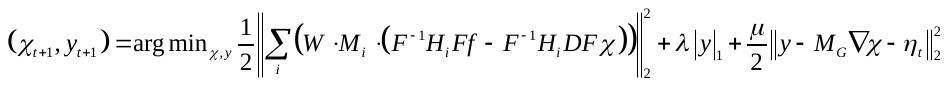
(11)

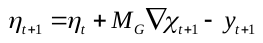
(12)

Here *μ* is a gradient consistency term parameter that has influence only on the convergence speed and not on the solution itself [14].

Equation (11) is minimized separately with respect to *χ* and *y*:

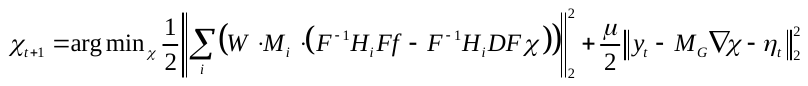
(13)

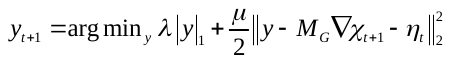
(14)

Equation (14) is solved by the soft-thresholding operation as proposed and implemented by Bilgic et al [14]:

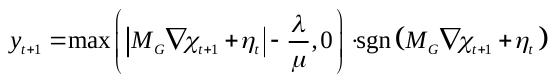
(15)

Equation (13) contains only *L*_2_ norms and its minimum can be found by setting its gradient to zero. Keeping in mind that *H_i_^H^* = *H_i_*, *D^H^* = *D* (H denotes the Hermitian conjugate transpose):

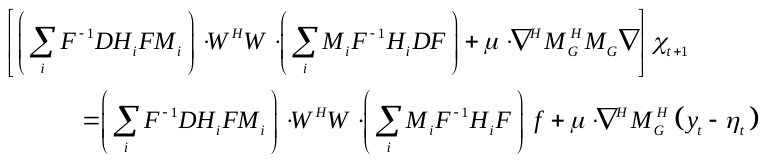
(16)

The gradient operator ∇ is calculated using a multiplication in k-space:

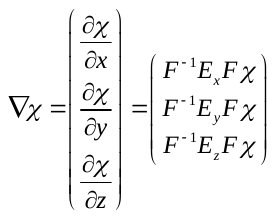
(17)

where *E_x_*, *E_y_* and *E_z_* are k-space representations of differentiation operators along image axes x, y and z. To reduce the number of Fourier transforms per operation, the minimization is performed relative to the susceptibility in *k*-space *Fχ*, and therefore the leftmost inverse Fourier and rightmost Fourier transform in both left and right sides of (16) are dropped:

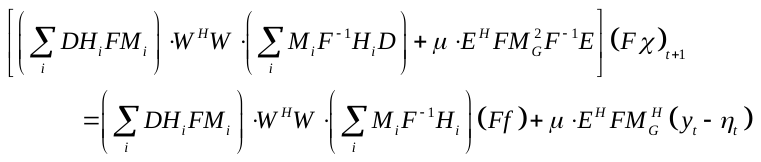
(18)

This equation is solved with the linear preconditioned conjugate gradient method with the following preconditioner, proposed in [8] to solve a *L*_2_-norm regularized QSM reconstruction problem employing multiple SMV kernels:

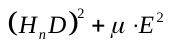
(19)

where *H_n_* is the discrete Fourier transform of the largest SMV kernel.
